# Building a single cell transcriptome-based coordinate system for cell ID with SURE

**DOI:** 10.1101/2024.11.13.623403

**Authors:** Feng Zeng, Jiahuai Han

## Abstract

The growing scale of single cell data demands standardized cell identification. We develop SURE, a data-driven method that establishes a cell coordinate system through optimally-sized metacells exhibiting enhanced transcriptional homogeneity. This approach enables precise cellular state characterization while providing three key atlas functionalities: zero-shot query-to-reference mapping, out-of-atlas detection, and cell type annotation. SURE’s hierarchical assembly pipeline successfully integrates large-scale atlas datasets, as demonstrated by constructing the Human Blood Metacell Atlas (HBMCA) from 5 million cells. Our method addresses the fundamental challenge of consistent cell positioning across diverse experimental contexts, facilitating robust comparative analyses between healthy and disease states.

## Background

Revealing and classifying the type and subtype of cells lie at the core of life sciences [1–4]. For centuries, defining the identity of a given cell (cell ID) has relied on discrete, concrete, and qualitative descriptors of features such as morphology and function. The advent of single-cell techniques has given rise to numerous cell atlas datasets, presenting an unprecedented opportunity for systematically defining cell IDs. Studies of these cell atlases have revealed that cell types and cell subtypes/states vary along a continuum, challenging the traditional notions of discrete cell identities. Furthermore, the single-cell measurement is highly sparse, and a variety of technical and biological noise can interfere with accurate characterization. There is no doubt that a unified, single cell transcriptome-based coordinate system for cell ID is timely needed, and a comprehensive characterization tool for cell populations has to be developed for constructing this reference system.

Currently, two strategies exist for building reference systems for transcriptome-based cell ID. The first relies on cell type annotations mostly obtained from bulk sequencing of purified cell populations of known types. The other involves cell manifolds [5]. Although cell manifolds are based on single cell sequencing data, the reference systems yielded by manifolds are inherently local and potentially differing from one another due to their constructions being made by different datasets. Both approaches are very valuable and had greatly facilitated research in the field. However, the former is constrained by the availability and accuracy of prior annotations and the latter has inherent limitations of unsupervised manifold learning techniques. Both of them are unable to fully meet the need for the establishment of a data-driven, unified reference system for cell ID in analyzing single cell transcriptomes.

In this work, we propose leveraging metacells to establish a coordinate system that provides a reference framework for cell ID. The concept of metacell was initially introduced to address the challenges of sparsity and noise in single-cell genomics data [6]. A metacell represents small, homogeneous groups of cells identified by the minimum unit of homogeneity. Employing metacells should make it feasible to construct a coordinate system that can serve as a reference for cell ID. First, since a set of metacells should represent the distribution of cell types and subtypes in a given single cell transcriptome dataset, metacells can serve as landmarks for mapping, aligning, and assembling different datasets. This lays the foundation for establishing a global, standardized coordinate system for all cell types. Notably, this process is data-driven, circumventing the restriction of prior annotation of known cell types. Second, metacells pool the information from homogeneous cells and reduce the impact of single-cell sequencing noise and sparsity, providing a more precise representation of cell ID. Third, metacells provide a compressed representation of cell atlas. By summarizing the high-dimensional single-cell data into a smaller set of metacells, metacells enable the efficient analysis of large-scale single-cell datasets, opening up new avenues for data exploration and hypothesis generation.

Several computational methods for metacell calling have evolved from cell clustering, employing discriminative approaches to partition cell data based on similarity, including Metacell [7], Metacell2 [8], SuperCell [9], and SEACells [10]. However, these clustering-based strategies are not well-suited for constructing a unified coordinate system. The objective of defining metacells for the coordinate system differs from metacells defined by cell clustering, as the former should capture the inherent continuity of cell states, transcending the discrete boundaries imposed by clustering algorithms. As a common problem, accurately defining cell similarity remains challenging due to the extreme sparsity inherent to single-cell genomics data, and existing metacell calling methods have not modeled and eliminated the impact of technical variations, especially batch effects, as well as the random differences among samples, rendering it difficult to project other single-cell data onto the coordinate system formed by these metacells. MCProj [11] attempts to achieve the projection of single-cell data by iteratively eliminating the differences between the query data and the metacell atlas, but this iterative algorithm struggles to meet the requirements of a unified coordinate system in terms of consistency, accuracy, and efficiency when dealing with large-scale, noisy data. Therefore, constructing a unified coordinate system requires new tools to overcome the limitations of existing metacell calling approaches.

In quest to define metacells for the coordinate system, we draw inspiration from vector quantization (VQ), a classic data compression technique from information theory [12]. VQ revolves around a codebook, a collection of codewords serving as data prototypes that capture the dataset’s characteristics with minimal information loss. The VQ-based generative approach diverges from the traditional cluster-based definition of metacells, conceptualizing the codebook as a collection of metacells, each representing a fundamental cell type or subtype. By employing VQ and deep generative modeling, we have achieved a principle for defining metacells to establish a coordinate system for cell ID. Leveraging the codebook’s compact representation, we discover metacells with enhanced representational power, comprehensively capturing the distribution of cell states and overcoming technical noise, i.e., batch effects. These metacells establish a unified coordinate system that accurately reveals salient features and underlying patterns, even when single-cell data quality is suboptimal. Metacells based on the VQ generative model effectively addresses cell type imbalance and enable the assembly of multiple single-cell atlas datasets through a hierarchical strategy. Using this divide-and-conquer approach, we have established a coordinate system for the cells from human hematopoietic system using single-cell transcriptome data. This metacell-based coordinate system supports the mapping of query data without requiring batch correction or model fine-tuning, making it a completely zero-shot process. Our method for establishing a coordinate system provides a technical foundation for constructing a unified reference for cell ID, paving the way for advancing single-cell data integration and analysis.

## Results

### A generative model for metacell calling

SURE (SUccinct REpresentation of cells) formulates metacell identification as a density estimation problem. This method conceptually places a uniformly distributed grid of metacells over the cell state space, where each metacell severs as a landmark characterizing local distributions. Using stochastic variational inference (SVI) [13], SURE continuously optimizes metacell positions to obtain precise estimates of cell state distributions and accurate reconstructions of single-cell transcriptional profiles (**Fig. 1a**).

**Fig. 1.**
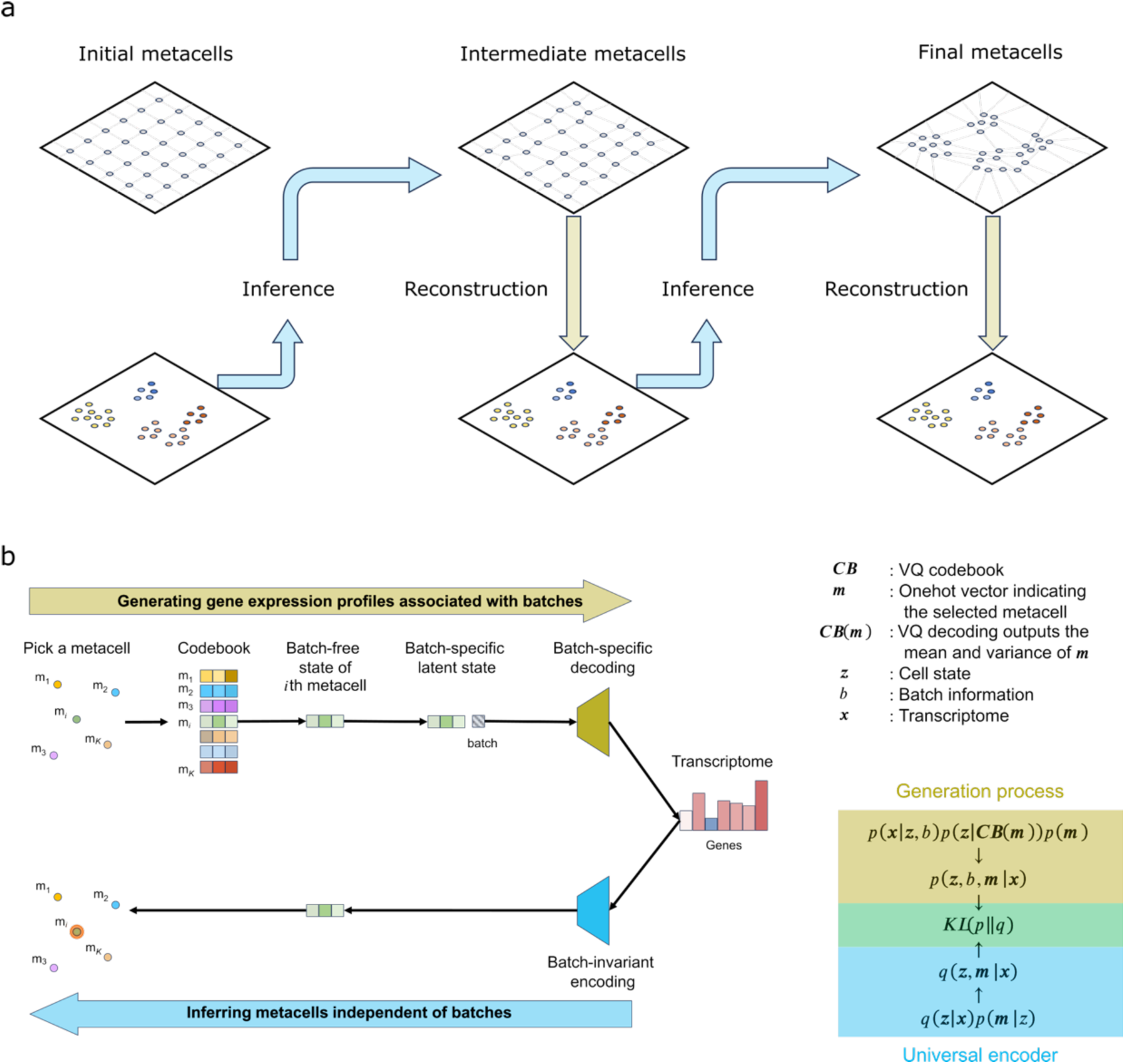
SURE model design. **a**, Illustration of SURE’s generative topographic mapping. **b**, SURE’s asymmetric variantional inference architecture.

This approach draws inspiration from self-organizing map (SOM) [14], an unsupervised neural network method that projects high-dimensional data onto a low-dimensional grid while preserving topological relationships through competitive learning. SURE also builds upon generative topographic mapping (GTM) [15], a probabilistic alternative to SOM that uses maximum likelihood estimation to position latent points in data space while maintaining a continuous mapping between latent and observed spaces.

SURE advances these approaches by integrating VQ with deep generative modeling (**Fig. 1b**). This framework introduces two key innovations: first, it employs a deep neural network to generate VQ codebook (composed of metacells), optimized through SVI rather than maximum likelihood estimation used in traditional approaches like SOM, GTM, and VQ-VAE [16]. Second, it implements a batch-invariant encoder through an asymmetric Bayesian architecture inspired by SCALEX, specifically designed to address the unprecedented technical variations (batch effects) prevelant in single-cell data that were not considered in SOM, GTM, and VQ-VAE. The VQ codebook’s metacells serve as batch-effect-free landmarks in a universal cell state space. During the generation of single-cell observations, batch-specific factors are incorporated to adapt these batch-invariant metacells to accommodate dataset-specific technical characteristics. This design enables the encoder to automatically identify and disentangle biological patterns (e.g., cell type or disease) from technical variations through SVI optimization, thereby facilitating zero-shot mapping of new datasets onto established cell atlases while maintaining biological fidelity.

To accommodate single-cell data complexity, SURE provides multiple modeling options:

1. Four latent variable distributions (Student-t, Gaussian, Laplacian, Cauchy);
2. Three count data distributions (multinomial, negative-binomial, Poisson).

Users can select appropriate distributions based on their specific data characteristics. This flexibility allows SURE to better capture the statistical properties of diverse single-cell datasets while maintaining computational efficiency through its variational inference framework.

### SURE identifies high-quality metacells

The generative modeling framework provides SURE with distinct advantages over alternative metacell calling methods. Using a single-individual scRNA-seq dataset containing CD34+ early hematopoietic cells [10], we evaluated SURE’s performance relative to alternative approaches. First, SURE produces metacells with more biologically reasonable size distributions. The number of cells assigned to a metacell (gamma parameter) significantly influences result quality, where excessively large metacells may obscure cell state boundaries while overly small metacells less effective at addressing single-cell data noise. SURE employs prior distribution modeling of latent variables to regulate metacell size, implementing four distribution types: Guassian, Student-t, Laplacian, and Cauchy. Anallysis of the CD34+ dataset showed that Laplacian and Cauchy priors generated both small and large metacells, Student-t prior permitted large metacells, while Gaussian prior restricted extreme sizes (**Fig. 2a**). Comparative evaluation revealed that alternative methods produced atypical size distributions: SEACells and SuperCell generated metacells with extreme sizes, MetaQ [17] failed to produce large metacells, and Metacell2 could not generate small metacells (**Fig. 2b**). These results demonstrate that SURE’s adaptive capability to identify appropriate metacell sizes that better match inherent data characteristics, where cell populations typically consist of dominant, intermediate, and rare cell states.

**Fig. 2.**
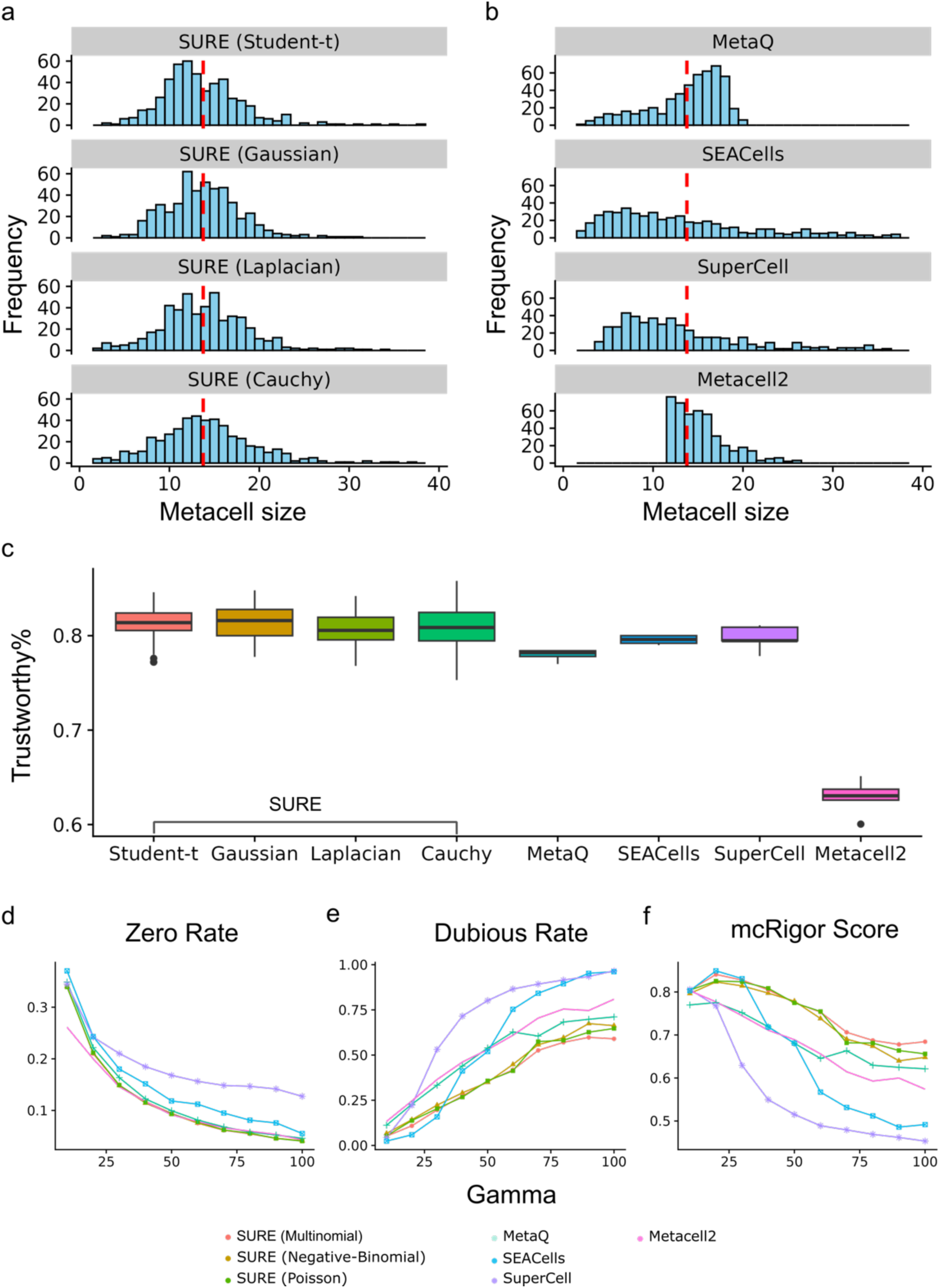
Metacell calling performance evaluation. **a-b**, Distributions of metacell sizes across different calling methods. The red line indicates the expected gamma value. **c**, Assessment of transcriptional homogeneity within individual metacells. **d-f**, Systematic evaluation of metacell performance as a function of size parameter gamma, including: (**d**) reduction of transcriptome sparsity (zero rate), (**e**) preservation of transcriptional homogeneity (dubious rate), and (**f**) overall performance scores derived from mcRigor analysis.

We applied mcRigor’s statistical framework to assess transcriptional homogeneity within metacells [18]. This method classifies metacells with homogeneous transcription profiles as trustworthy and those with divergent profiles as dubious. Our evaluation showed that Student’s t prior yielded more trustworthy metacells than other priors, and SURE consistently outperformed alternative methods in producing trustworthy metacells (**Fig. 2c**).

Through systematic variation of gamma parameters using mcRigor’s optimization functionality, we calculated two key metrics: dubious rate (DubRate, proportion of cells in questionable metacells) and zero rate (ZeroRate, transcriptome sparsity in aggregated metacells). The results demonstrated SURE’s effectiveness in addressing sparsity while controlling dubious rates across gamma values (**Figs. 2d** and **e**). At low gamma values, both SURE and SEACells achieved lower dubious rates than MetaCell2, SuperCell, and MetaQ, with SURE uniquely maintaining low zero rates. In contrast, SEACells and SuperCell metacells retained substantial zero values in their aggregated transcriptomes. As gamma increased, SURE’s advantages became more pronounced, consistently producing metacells with minimal dubious and zero rates, while alternative methods showed significant increases in questionable metacells. Notably, even at high gamma values, SEACells and SuperCell continued to exhibit zero values in metacell transcriptome, further demonstrating the effectiveness of SURE’s likelihood approach in generating reliable metacells with low-sparsity transcriptional profiles across parameter configurations.

Regarding UMI count modeling, SURE provides multiple distribution options (negative-binomial, multinomial, and Poisson, with zero-inflation options) to accommodate single-cell data complexity. Benchmarking revealed no single distribution demonstrated clear superiority, through multinomial distribution achieved higher mcRigor scores at low gamma values. No significant performance differences were observed between zero-inflation and standard models (**Figure S1**).

### SURE enables calling consensus metacells across samples

Current metacell calling methods face challenges in deriving consensus metacells from multiple samples. SEACells employs a two-step approach involving initial metacells construction for individual samples followed by batch correction (e.g., using Harmony) and subsequent consensus metacell identification. However, as demonstrated previously, such methods inevitably introduce distortions into aggregated transcriptomes. SURE offers a more effective solution for direct consensus metacell detection from multiple samples.

For evaluation, we utilized the healthy human BMMC dataset provided by the NeurIPS 2021 Open Problems in Single-cell Analysis competition [19] (n=69,249 cells from 13 batches, generated using 10X Multiome technology, **Fig. 3a**). While the dataset includes both RNA and chromatin accessibility data, our analysis focused exclusively on RNA data. We compared SURE, MetaQ, and SEACells in identifying consensus metacells across 13 batches, with UMAP visualization showing SURE’s metacell distribution more closely resembled the original single-cell distribution (**Figs. 3b-d**).

**Fig. 3.**
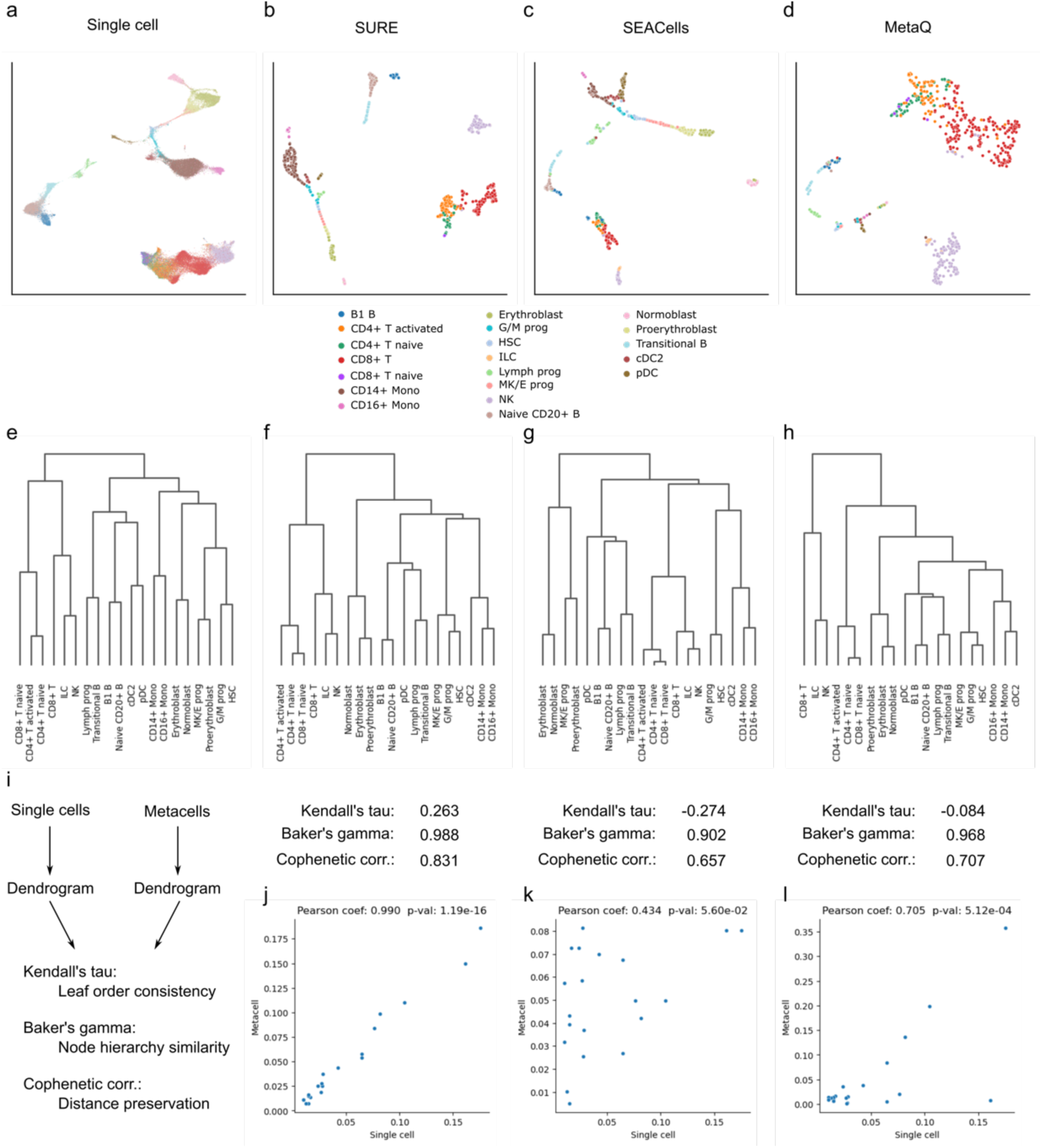
Evaluation of metacell identification in multi-batch datasets. **a-d**, UMAP visualizations comparing cell population distributions and metacell identification results between SURE and alternative methods. **e**, Reference cell lineage tree constructed from single-cell transcriptional profiles. **f-h**, Lineage trees reconstructed from metacell transcriptomes generated by SURE (**f**) and alternative methods (SEACells (**g**) and MetaQ (**h**)). **i**, Quantitative comparion of lineage tree topologies between metacell-derived reconstructions (from left to right: SURE, SEACess, and MetaQ) and the reference tree. **j-l**, Correlation analyses of cell type frequencies between metacell-derived and original single-cell data for SURE (**j**), SEACells (**k**), and MetaQ (**l**).

To assess the biological fidelity, we constructed cell lineage trees from consensus metacells and compared them to single-cell-derived cell lineage using quantitative metrics (**Figs. 3e-i**):

1. Kendall’s tau (measuring the agreement in pairwise ordering relationships between leaves of two trees, where a value of 1 indicates perfect concordance in terminal node ordering) [20];
2. Baker’s gamma (quantifying the topological similarity in branching patterns between hierarchical tree structures by comparing the relative positions of internal nodes) [21];
3. Cophenetic correlation (assessing the preservation of pairwise phylogenetic distances between corresponding nodes in different tree representations) [22].

Resuts demonstrate that SURE’s metacells accurately preserved the original cell lineage structure, while MetaQ and SEACells introduced significant distortions. Furthermore, SURE better maintained cell population compositions, as evidenced by Pearson correlation analysis between single-cell and metacell state frequencies (SURE: Pearson coef.= 0.990, p-value: 1.19e-16; SEACells: Pearson coef.=0.434, p-value: 5.60e-2; MetaQ: Pearson coef.=0.705, p-value:5.12e-4, **Fig. 3j-l**). These findings collectively indicate SURE’s superior performance in consensus metacell identification while preserving biological structures.

### Leveraging metacell codebook to augment cell atlas functionality

Global efforts in cell atlas construction are ongoing. However, limitations in atlas summarization, query, and retrieval methods constrain the functionality of cell atlases. SURE demonstrates several advantages that make it effective for atlas management and reuse (**Fig. 4a**).

**Fig. 4.**
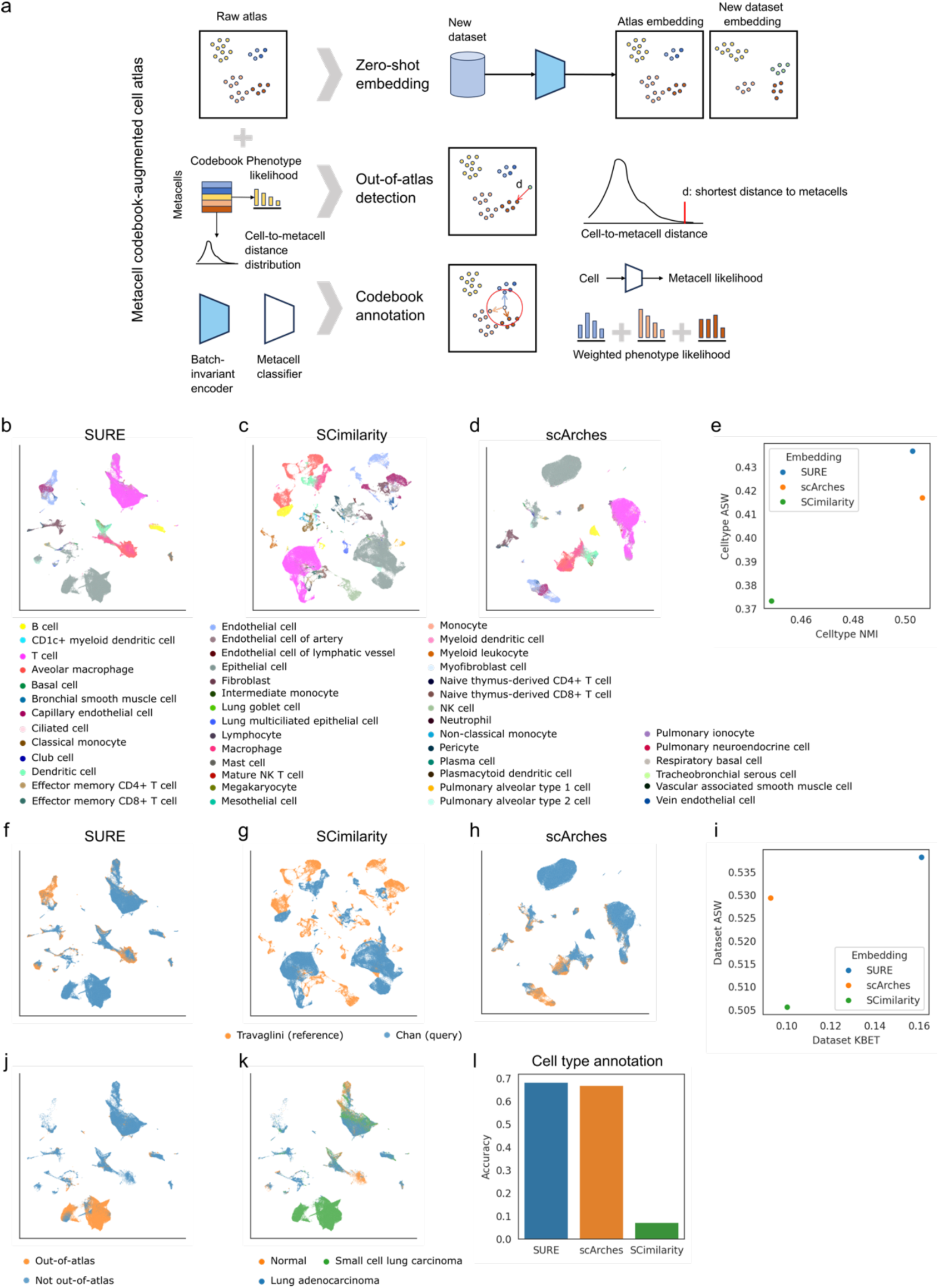
Codebook-augmented cell atlas analysis. **a**, Schematic overview of metacell codebook functionalities, including zero-shot mapping, out-of-atlas detection, and phenotype annotation capabilities. **b-d**, UMAP visualizations comparing cell type distributions in query-to-reference mapping results produced by SURE, SCimilarity, and scArches. **e**, Quantitative assessment of cell type preservation performance across mapping results. **f-g**, UMAP projections demonstrating data integration quality for reference and query datasets processed by SURE, SCimilarity, and scArches. **i**, Systematic evaluation of dataset merging characteristics in query-to-reference mapping. **j**, Detection results of out-of-atlas cell populations in query data. **k**, Disease state annotations of query cells. **l**, Performance metrics of cell type annotation accuracy.

First, SURE’s batch-invariant encoder enables the zero-shot mapping of new datasets onto the atlas. This eliminates the need for fine-tuning, which typically requires considerable computational time. To evaluate zero-shot mapping capability, we used the Travaglini dataset [23] of normal human lung cells as the reference. We compared SURE with SCimilarity (a foundation model supporting zero-shot mapping) [24] and scArches (which employs a fine-tuning strategy) [25]. The query data consisted of the Chan dataset [26] containing healthy and malignant cells. Both SURE and SCimilarity embedded query cells into the reference space without additional learning, while scArches required fine-tuning (**Figs. 4b-d**). Evaluation using average Silhouette width (ASW) and normalized mutual information (NMI) revealed that SURE achieved superior cell type cohesion (cell type ASW=0.437) compared to scArches (cell type ASW=0.417) and SCimilarity (cell type ASW=0.373) (**Fig. 4e**). Furthermore, SURE demonstrated excellent dataset mixing performance (dataset ASW=0.538, KBET=0.161), outperforming both SCimilarity (dataset ASW=0.506, KBET=0.100) and scArches (dataset ASW=0.529, KBET=0.093) (**Figs. 4f-i**).

Second, SURE facilitates detection of query cells outside the atlas distribution. During the atlas construction, SURE calculates cell-to-metacell distances to establish a background distribution model. This enables the identification of out-of-atlas cells exhibiting either biologically significant characteristics or extreme batch effects. Application to the Chan dataset (p-value < 0.01) revealed that most out-of-atlas cells were epithelial cells specific to small cell lung carcinoma (**Figs. 4j** and **k**).

Third, SURE enables precise cell type annotation through metacell codebook querying. The codebook not only encodes the latent states of metacells but stores phenotype information summarized from individual cells, i.e., the cell type likelihood. SURE calculates metacell likelihoods indicating cell-to-metacell associations. Aggregation of the codebook and metacell likelihoods produce cell type likelihoods for individual cells. Annotation occurs via maximum likelihood estimation (MLE). Comparative evaluation using the Chan dataset showed that SURE’s annotation accuracy (0.684) exceeded scArches (0.670) and SCimilarity (0.073) (**Fig. 4l**).

### SURE enables cell atlas assembly

The construction of universal cell atlases presents significant computational challenges due to the intensive processing requirements and demanding hardware specifications. SURE addresses these limitations through metacell-based atlas compression, where the substantially reduced number of metacells (relative to the original atlas size) enables efficient data integration (**Fig. 5a**). SURE implements a bootstrapping strategy that generates representative subsets of pseudo-cells while preserving transcriptional profiles (**Fig. 5b**). This process involves: (1) sampling from each metacell’s local distribution, (2) identifying nearest-neighbor cells to these samples through spatial search, and (3) aggregating transcriptional profiles from sampled cells. Repeated iterations produce bootstrap-sampled pseudo-cells suitable for universal codebook construction across multiple atlas datasets.

**Fig. 5.**
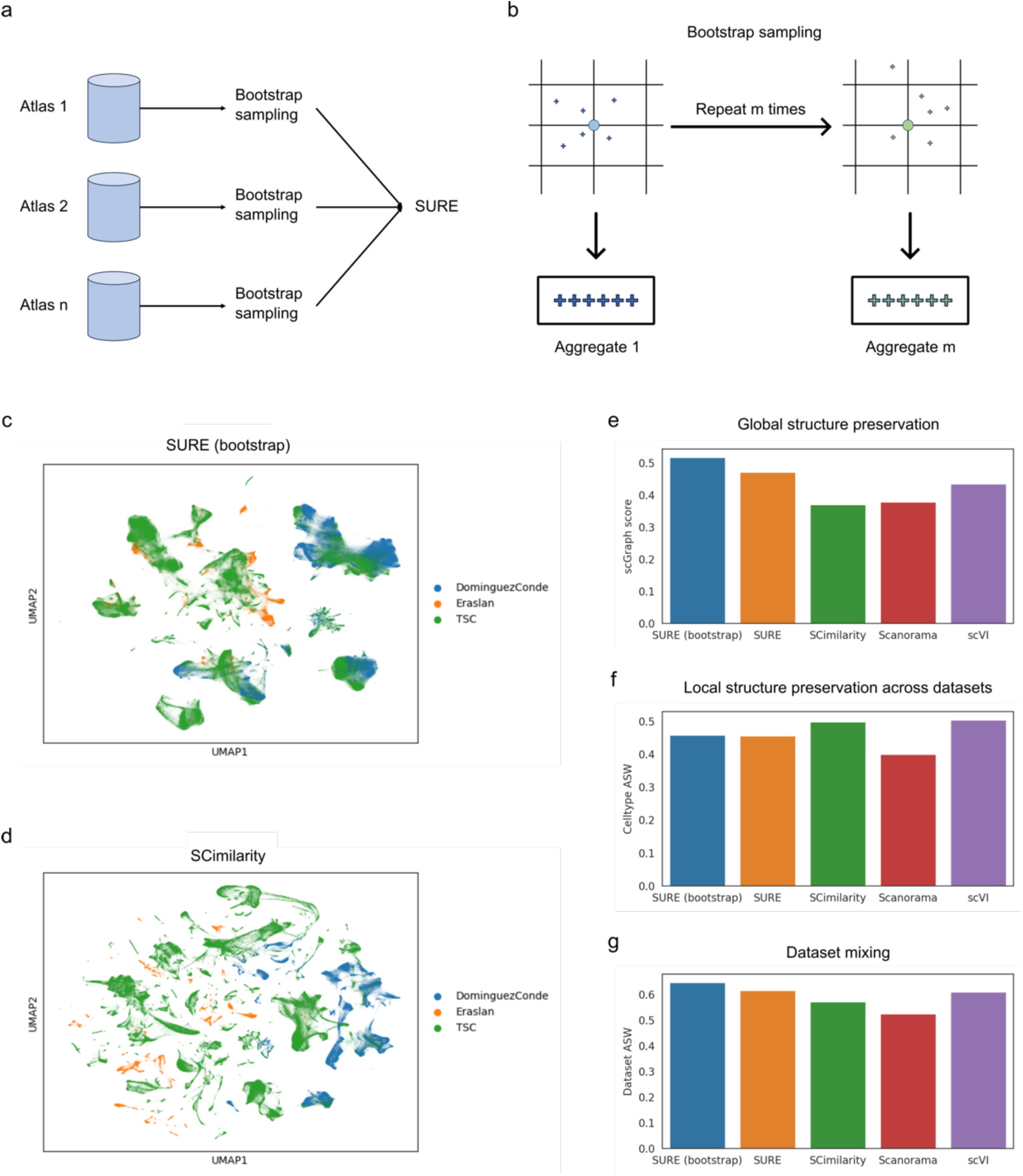
Hierarchical cell atlas assembly framework. **a,** Schematic representation of the hierarchical cell atlas assembly pipeline implemented in SURE. **b**, Bootstrap sampling methodology based on cell state space density estimation derived from SURE analysis. Circle points are metacells. Cross points are sampled cells. **c-d**, Comparative UMAP visualizations of large-scale atlas integration results generated by SURE and SCimilarity, demonstrating spatial organization of integrated cell populations. **e-g**, Comprehensive evaluation metrics addressing: (**e**) preservation of global cellular neighborhood relationships, (**f**) maintenance of local cell type structures, and (**g**) effectiveness in mixing heterogeneous atlas-scale datasets.

We evaluated SURE’s assembly performance against SCimilarity, Scanorama [27], and scVI [28] using three heterogeneous atlas datasets (DominguezConde [29], Eraslan [30], and TSC [31]) encompassing diverse cell types across multiple tissues. Two SURE variants were compared: metacell integration (SURE) and bootstrap-sampled pseudo-cell integration (SURE (bootstrap)). UMAP visualization demonstrated superior dataset mixing with SURE (bootstrap) compared to SCimilarity (**Figs. 5c** and **d**). Quantitative assessment using scGraph [32] revealed that SURE (bootstrap) better preserved global population structures than both the standard SURE implementation and alternative methods (**Fig. 5e**). The scGraph score assesses the preservation of cell-cell neighborhood relationships between atlas assembly results and their original single-cell datasets, measuring how faithfully biological structures are maintained during the assembly process.

Local structure preservation analysis yielded nuanced results: while SURE variants showed marginally lower cell type ASW scores than SCimilarity and scVI, they achieved superior dataset ASW scores (**Figs. 5f** and **g**). This pattern suggests SURE’s enhanced capability to integrate biologically equivalent cell types across different datasets despite potential annotation discrepancies (e.g., identical cell types with different nomenclature), where comparator methods tended to maintain these as distinct populations. These findings collectively demonstrate SURE’s effectiveness in assembling atlas datasets with varying cellular compositions, with bootstrap sampling further improving integration performance.

### HBMCA: Human Blood MetaCell Atlas

As an application, we utilized SURE to establish the Human Blood MetaCell Atlas (HBMCA), a cell coordinate system for the human hematopoietic system. We collected 29 public datasets, covering different types of cell suspensions (i.e., single cells and single nucleus) processed by various single-cell sequencing technologies, including 10x 5’ (v1, v2), 10x 3’ (v2, v3), Smart-seq2, Seq-Well, Microwell-seq, BD Rhapsody, and ScaleBio. This comprehensive dataset encompasses 1,906 individuals, with a total of 5,310,750 blood cells sequenced (**Fig. 6a**).

**Fig. 6.**
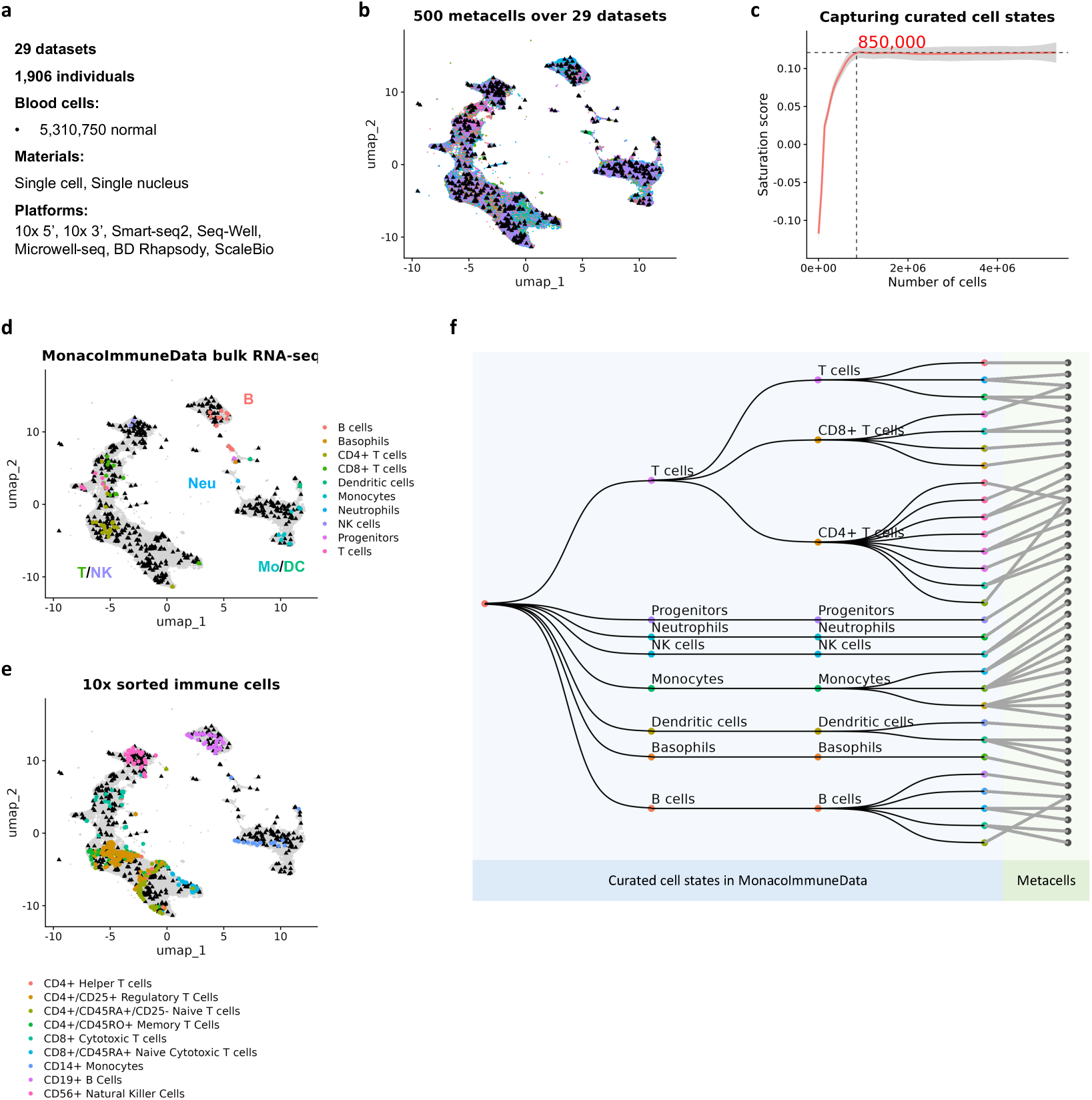
Human blood metacell atlas (HBMCA) characterization. **a**, Technical statistics of the integrated dataset. **b**, Visualization of the blood cell coordinate system constructed from metacells, with black triangles denoting individual metacell positions. **c,** Saturation analysis evaluating the coordinate system’s capacity to represent distinct cell populations, employing Silhouette scores to simultaneously quantify both intra-cell-type homogeneity and inter-cell-type separation. **d-e,** Validation through orthogonal datasets: (d) Projection of immune cell states identified through fluorescence-activated cell sorting and bulk sequencing onto HBMCA; (**e**) Mapping of single-cell sorted immune cells processed via 10x Genomics onto the atlas. Both projections use color-coding to distinguish established cell types. **f**, Systematic alignment of curated cell states from the MonacoImmuneData reference database with HBMCA metacells, demonstrating comprehensive coverage of known immulogical states.

Using SURE, we constructed a reference cell coordinate system comprising 500 metacells derived from five million blood cells, optimized through comprehensive evaluation of cell type purity, coverage, and pre-training complexity (**Figures S3-S5**). **Figure 6b** demonstrates this system’s effectiveness in integrating cell states across diverse 29 datasets. To determine the minimal number of cells required for representative performance, we conducted a saturation analysis by iteratively building coordinate systems from progressively larger cell subsets (ranging from 10,000 cells up to the complete dataset). Each system was evaluated by projecting all five million cells and calculating Silhouette scores to assess both intra-cell-type compactness and inter-cell-type separation. This analysis revealed that the system’s representational capacity stabilized when constructed from more than 850,000 cells, empirically validating that our five million cell-based coordinate system achieves comprehensive coverage of blood cell type heterogeneity.

To further analyze the similarities and differences between the metacell-based human blood cell type atlas and existing reference databases for blood cell types, we performed comparisons using MonocoImmuneData and ImmGenData [33]. The former utilized cell sorting techniques to isolate 29 cell types, while the latter isolated 20 cell types, both employing bulk RNA sequencing. Additionally, we utilized single-cell data of sorted immune cells provided by 10x Genomics[34]. We mapped these reference databases onto HBMCA through SURE (**Figs. 6d** and **e**, **Figures S6-S8**). Based on these mapping results, we observed that the four major cell populations in HBMCA corresponded to T/NK cells, B cells, monocytes, and neutrophils in these databases. The verification using bulk RNA sequencing data and single-cell sequencing data demonstrated the accuracy and reliability of our established cell coordinate system. Furthermore, this result also indicated that HBMCA supports the mapping of not only single-cell data but also other types of transcriptomic data (i.e., bulk RNA sequencing).

Simultaneously, we observed that the data-driven HBMCA exhibits clear advantages over traditional cell type reference systems in delineating cell types. On one hand, HBMCA provides a more comprehensive characterization of cell types. Traditional cell type reference systems can only capture a small portion of the blood cell atlas. As depicted in **Fig. 6f**, the 29 sorted human immune cells from MonocoImmuneData mapped to 43 metacells, and based on the proportion of metacells, they are estimated to cover 8.6% of the blood cell atlas. On the other hand, HBMCA offers a more refined characterization of cell types. The resolution of traditional reference systems is limited; among the 29 sorted human immune cells, 10 (approximately 34.5%) mapped to 2 metacells, and 3 (approximately 10.3%) mapped to 3 metacells, respectively. Therefore, the data-driven cell coordinate system possesses a more comprehensive ability to define cells.

### HBMCA enables the visualization of variations across studies and illnesses

The establishment of HBMCA represents a significant milestone in single-cell research, enabling the comparative analysis of single-cell data from diverse studies and diseases within a unified reference coordinate system. HBMCA boasts several notable advantages. Firstly, it stands as the most comprehensive human blood single-cell atlas database to date, providing a characterization of the distribution of cell states in the normal human hematopoietic system (**Fig. 7a**). Secondly, HBMCA’s zero-shot capability allows for the direct projection of single-cell data from other studies onto its reference framework, eliminating the need for batch correction or model fine-tuning. This feature renders HBMCA an invaluable resource for investigating the impact of various diseases on the hematopoietic system.

**Fig. 7.**
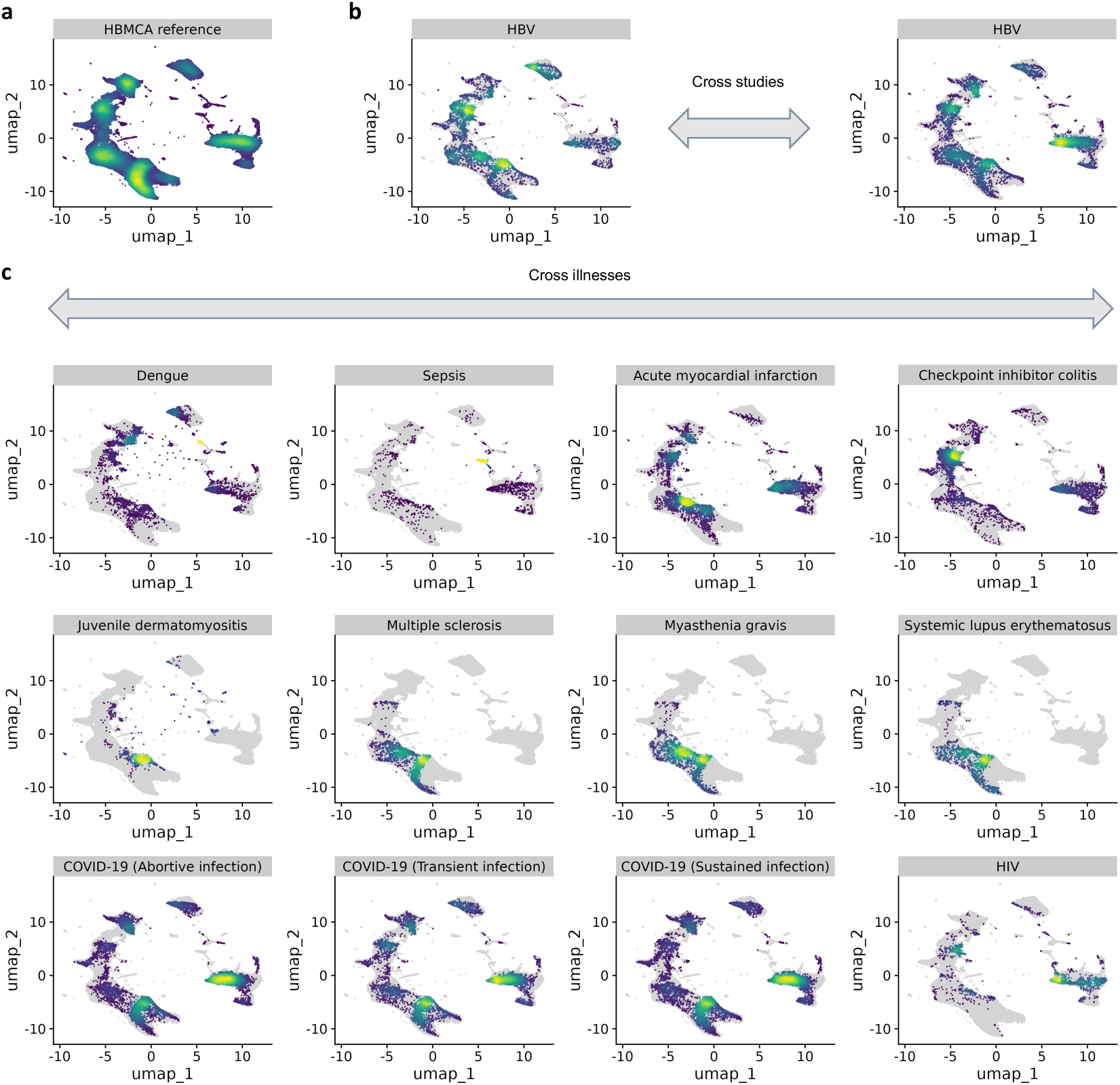
Zero-shot mapping of cells across studies and illnesses onto HBMCA. **a**, Distribution of cells collected from normal individuals. **b**, Illustration of cell distributions across two HBV studies. **c**, Illustration of cell distributions across twelve disease conditions.

To showcase this functionality, we employed two single-cell datasets focusing on hepatitis B virus (HBV) infection (**Fig. 7b**). The first dataset comprised single cells primarily derived from Asian HBV patients[35], while the second dataset contained single-cell data from European HBV patients[36]. These two studies were conducted independently by different research teams using distinct single-cell technology platforms. By leveraging SURE’s universal encoder, we successfully projected these two HBV datasets onto HBMCA. The results unequivocally demonstrated that the single-cell data from these independent studies revealed remarkably similar effects of HBV on the hematopoietic system, with their single-cell findings exhibiting high reproducibility and corroboration.

To further illustrate the versatility of HBMCA, we curated single-cell data from 12 different illnesses, encompassing a wide spectrum of conditions such as dengue virus infection [37], sepsis [38], acute myocardial infarction [39], checkpoint inhibitor-induced colitis [40], juvenile dermatomyositis [41], multiple sclerosis [42], myasthenia gravis [42], systemic lupus erythematosus [42], various types of SARS-CoV-2 infection (including abortive, transient, and sustained infection) [43], and HIV infection [44] (**Fig. 7c**). Utilizing SURE, we directly projected these datasets onto HBMCA, circumventing the need for batch correction or model fine-tuning. Within the unified coordinate system provided by HBMCA, we were able to observe and analyze the changes in cell states of the human hematopoietic system under these diverse disease conditions.

## Discussion

The establishment of standardized coordinate systems has played a pivotal role in genomic research by enabling direct mapping and comparison of genomic data across different laboratories. Similarly, single-cell omics research urgently requires a unified coordinate system to facilitate the integration and comparison of diverse cell atlas datasets. In this study, we propose the use of metacells as fundamental units for constructing such a coordinate system and develop SURE, a novel framework that employs vector quantization and deep generative modeling to learn metacell representation from single-cell transcriptome data.

Our analyses demonstrate that SURE offers several key advantages over existing approaches. First, SURE’s metacells more accurately capture the underlying cell-state distributions in atlas datasets. While current metacell identification methods often produce metacells with extreme size distributions that compromise their representativeness, SURE’s probabilistic framework generates metacells with higher transcriptional homogeneity and better resolves data sparsity issues, as confirmed by mcRigor’s statistical testing. This enhanced representation capability stems from SURE’s transcriptome-wide generative modeling approach that leverages count data likelihood.

Second, SURE introduces an innovative asymmetric Bayesian inference method for direct identification of consensus metacells across batches, outperforming the two-step process used in SEACells. Through comprehensive lineage comparative analysis using three distinct tree metrics, we show that SURE’s joint approach to metacell identification and batch correction better preserves cell population relationships. Frequency analyses further confirm that SURE maintains more accurate cell population compositions compared to methods that perform these steps separately, which can amplify distortions through sequential processing.

Third, the metacell codebook serves as an efficient compressed representation for large-scale atlas datasets. As single-cell techniques become more accessible and atlas datasets grow exponentially, SURE’s compressive representation and bootstrapping approach provide a scalable solution for atlas integration while preserving biological relationships and addressing technical variations (e.g., our HBMCA analysis shows that approximately 500 metacells can effectively represent over 5 million cells with a compression ratio up to 10,000:1).

Finally, SURE’s codebook empowers cell atlas functionality by enabling zero-shot mapping capability that outperforms alternative methods like SCimilarity and scArches. The framework incorporates statistical testing for detecting out-of-atlas cell populations and implements a codebook-based annotation strategy that accurately predicts cellular phenotypes through metacell-weighted voting, demonstrating superior performance compared to both contrastive learning and transfer learning approaches.

In conclusion, SURE establishes a robust coordinate system for cell identities using single-cell transcriptome data, with metacells serving as reference landmarks. The framework demonstrates promising performance in dataset harmonization, scalability, and consistent cell positioning across diverse datasets. This advancement not only facilitates more accurate cell atlas construction and integration but should also provide a foundation for elucidating regulatory networks underlying cell phenotypes, potentially contributing to the development of improved diagnostic and therapeutic strategies.

## Methods

### SURE

SURE provides an efficient method for integrating various single-cell datasets into a comprehensive cell atlas while defining a set of standard cell states or “metacells”. These metacells serve as robust reference landmarks, constituting a unified cell coordinate system for consistent cell positioning and annotations across a variety of datasets. SURE offers remarkable computational efficiency, enabling scalable analyses of large-scale single-cell data. Furthermore, metacells assure robust representations of cell diversity and effectively counteract data imbalance. Also, by leveraging the unified coordinate system, SURE systematically eliminates technical noises like batch effects, simplifying data integration and enabling meaningful biological comparisons across multiple datasets.

The underpinning of SURE lies in a variational Bayesian framework, whereby it integrates neural vector quantization and generative modeling techniques. These crucial aspects will be addressed in the following sections.

### Asymmetric Variational Bayesian Framework

Bayesian analysis forms an effective strategy for uncovering the true cell states in single-cell data. According to our model, the expression data of a cell is *x*, batch information is *b*, and its cell state is *z*. SURE’s objective is to create a unified cell coordinate system for the unified cell state space. Given the data *x*, SURE can accurately deduce the cell state *z* in a unified context through the inference model *q*(*z*|*x*). Notably, the omisson of batch information *b* in *q*(*z*|*x*) significantly widens the method’s applicability, indicating that, once adequately trained, the model can be universally deployed on new datasets without the necessity of batch information. Consequently, the inference of cell states is made batch-independent, aligning *q*(*z*|*x*) with *p*(*z*|*x*, *b*), and ensuring consistent, robust identification of cell types across diverse datasets.

The quest to realize a universal *q*(*z*|*x*) methodology presents considerable challenges. Single-cell data, being continuous and high-dimensional, results in an exponentially vast solution space for *z*. Additionally, the high sparsity of the data complicates inference, and batch effects often obscure true biological distinctions. These challenges are tackled by borrowing insights from the scVI’s generative model, implementing the Bayesian equation:

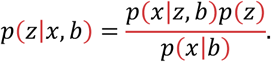

This equation provides a robust framework for deciphering *z* by contemplating the observed data *x* and batch information *b*, then going through the forward modeling of data generation *p*(*x*|*z*, *b*).

To successfully establish a universal *q*(*z*|*x*), we employ it as a variational approximation of *p*(*z*|*x*, *b*). This adaptation separates the inference and the generation models into two individual components: the generation model processes the cell state *z* and batch information *b* to produce *x*, thereby estimating *p*(*z*|*x*, *b*), while the inference model *q*(*z*|*x*) aims to approximate *p*(*z*|*x*, *b*) as closely as possible, i.e., *q*(*z*|*x*) ≈ *p*(*z*|*x*, *b*). We propose that this approximation is validated when the training data provide sufficient information about batch variations. Fortunately, over the past decade, tremendous amounts of data have been generated globally through various single-cell sequencing technologies, covering all possible batch variations.

Thus, SURE exhibits a distinct and asymmetric structure. While batch information *b* remains a deterministic variable in the generation process, SURE trains the inference model *q*(*z*|*x*) using purely *x*, thereby sidelining batch information and reinforcing the model’s universal applicability.

### Neural Vector Quantization

To address the high-dimensional, continuous nature of the cell state *z*, we implement neural vector quantization to reduce solution space complexity. We propose a codebook containing *K* landmarks within the cell state space *Z* = {*z*}, designed to provide comprehensive coverage of *Z* with appropriate resolution. Each cell’s state *z* can then be characterized by its proximity to the nearest landmark *i* through the probabilistic model *p*(*i*)*p*(*z*|*i*):

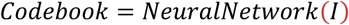

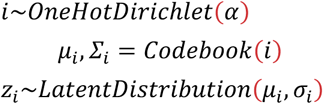

Here, *I* is a unit matrix indexing all landmarks, while *α* denotes a prior distribution over *K* landmarks. In our implementation, *α* uses a noninformative prior, assuming initial metacells are uniformly distributed in the cell state space. SURE employs a multi-layer neural network functions as the codebook for vector quantization, capable of outputting parameters for the latent distribution associated with each landmark. The framework supports four latent distribution types (Student-t, Gaussian, Laplacian, and Cauchy), all belonging to the local-scale distribution family. Unlike Gaussian distributions, the Student-t, Laplacian, and Cauchy distributions accommodate long-tail phenomena, making them particularly suitable for handling extreme in single-cell data (e.g., large cell populations or rare cell states).

The Vector Quantization Variational Autoencoder (VQ-VAE) provides a foundational approach for discrete latent representation learning. In VQ-VAE:

1. Continuous latent variables are replaced with discrete codes from a fixed codebook.
2. The Expectation-Maximization (EM) algorithm performs maximum likelihood estimation to learn latent discrete variables.
3. Each data point is assigned to the nearest codebook vector through hard assignment.
4. The original data is approximated by discrete metacells.

While VQ-VAE represents a classical approach using discrete points to model latent space, our approach differs in several key aspects:

1. Our codebook employs a dynamic multi-layer neural network rather a static data structure, providing greater flexibility and representational capacity.
2. Instead of discrete point representations, our neural network-based codebook describes data probability density through a superposition of *K* discrete latent distributions centered at these landmarks. SURE uses metacells to define the latent distributions accounting for the original transcriptional observations.
3. We utilize Bayesian inference for codebook learning, as opposed to the EM algorithm typically employed in VQ-VAE, enhancing our model’s ability to capture complex data structures within a probabilistically coherent framework.

### Generative Modeling

By integrating the Bayesian framework with neural vector quantization, we formulate SURE’s generative model through the following probabilistic process:

1. **Metacell selection:** A metacell (codebook element) is randomly selected as the initial state *i* for each cell, following by the prior distribution *p*(*i*).
2. **Biological state generation:** The batch-effect-free cellular state *z*is sampled from the probability distribution *p*(*z*|*i*) associated with the selected metacell.
3. **Batch effect incorporation:** The batch-specific cellular state [*z*, *b*] is generated by combing the the biological state *z* with batch factor *b*.
4. **Observation generation:** The observed cellular data *x* is generated from the batch-aware cellular state [*z*, *b*] through a count distribution model *p*(*x*|*z*, *b*). SURE implements three core count distributions (multinomial, negative-binomial, and Poisson), each with optional zero-inflation extensions to handle dropout events.

This generative process can be formally expressed in the joint probabilistic distribution:

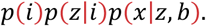

The corresponding variational approximation model is defined as:

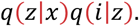

For implementation, we employ the Pyro probabilistic programming language to construct both the generative and variational models. Parameter optimization is achieved through stochastic variational inference, which provides scalable learning for large single-cell datasets while maintaining theoretical guarantees.

### Zero-shot cell mapping and UMAP computation

The asymmetric Bayesian inference framework has been demonstrated effectiveness in learning biological patterns from single-cell data while mitigating technical noise. Building upon this architecture, SURE’s encoder *q*(*z*|*x*) enables zero-shot mapping capability that directly projects new datasets into the learned latent space without requiring additional learning steps such as fine-tuning or transfer learning. This functionality enhances the utility of cell atlases by allowing rapid assessment of cell population compositions in newly acquired datasets while identifying novel cell states or unexpected technical variations. The architecture’s asymmetric design ensures stable performance across datasets with minor variations, providing biologically meaningful representations that maintain consistency with reference atlases.

The latent representations generated by *q*(*z*|*x*) can be directly visualized using UMAP. However, for query-to-reference mapping applications involving new datasets, performing joint UMAP embedding of both reference and query data requires substantial computational resources. To address this limitation, we implement an efficient visualization pipeline where a UMAP model is pre-trained exclusively on the reference atlas. This pre-trained model is then applied directly to project query datasets into the established visualization space, significantly reducing computational overhead while maintaining biological interpretability.

### Metacell classification and phenotype classification

SURE’s architecture enables direct classification of new single-cell data points to existing metacells through its dual encoding framework. The process first maps input data into latent space using the encoder *q*(*z*|*x*), then assigns cells to metacells through the classification function *q*(*i*|*z*). This two-stage approach distinguishes SURE from alternative metacell calling methods, which lack this integrated classification capability and cannot directly assign new data points to established metacell structures.

The metacell classification probabilities derived from *q*(*i*|*z*) enable the construction of a phenotype landscape through aggregation of individual cell characteristics. This landscape can represent various phenotypic attributes including cell type annotations, disease states, or experimental conditions.

Formally, let *P* ∈ {0,1}^*n*×*k*^ denote the phenotype identity matrix for *n* cells across *k* phenotype categories (e.g., distinct cell types/states), where each row is a one-hot encoded vector indicating the specific phenotype of each cell. Let *Q* ∈ [0,1]^*n*×*m*^ represent the probabilistic metacell assignment matrix for *n* cells across *m* metacells, where row sums are normalized to unity.

The phenotype potential landscape for metacells is computed as:

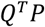

followed by row-wise normalization. This computationally efficient representation allows SURE to perform rapid phenotype annotation of new dataset through metacell-based weighted classification based on the probabilistic cell-to-metacell assignments.

### Out-of-atlas detection

SURE implements a robust statistical framework for identifying novel cell states and technical variations in newly analyzed datasets. The methodology operates by first establishing a reference null distribution through Gaussian kernel density estimation of cell-to-nearest-metacell distances computed from the original atlas data in latent space. For query datasets, each cell’s distance to its closest metacells is calculated and evaluated against this reference null distribution. Cells demonstrating statistically significant deviation (p-value<0.01) are classified as potential out-of-atlas observations. These outliers are interpreted contextually: when predominantly representing isolated variations, they may indicate novel cell states; when query dataset exhibits systematic variation patterns, they likely reflect technical variations necessitating batch effect correction.

### Bootstrap sampling

SURE employs a bootstrapping methodology to sample from atlas datasets while preserving the distributional properties encoded in the associated codebook. The sampling procedure operates through an iterative process where, at each iteration:

1. **Latent space sampling:** k points are generated in the latent space following the probability distribution modeled by the codebook.
2. **Real data mapping:** For each generated point, the algorithm identifies the nearest neighboring real data points in the original dataset.
3. **Profile aggregation:** The transcriptional profiles of these selected real data points are aggregated to produce representative profiles.

This bootstrapping approach offers two key advantages over direct sampling from the raw atlas datasets:

1. **Noise reduction:** The aggregation process naturally smooths technical noise while preserving biological signals.
2. **Granularity preservation:** The resulting profiles maintain single-cell resolution despite the aggregation step.

The method effectively bridges the gap between metacell-level analysis and single-cell resolution by combining the statistical power of aggregated profiles with the precision of single-cell data representation.

### Cell atlas assembly

To address the challenges of large-scale cell atlas integration, we introduce a hierarchical assembly approach that systematically processes datasets through multiple resolution levels. This method begins by applying SURE to each individual dataset to generate primary metacells, which serve as the compressed yet biologically meaningful representations of the original single-cell data. These primary metacells capture dataset-specific biological variations while significantly reducing computational complexity.

Building upoin this foundation, the method proceeds to create bootstrapped samples from the primary metacells, a process that maintains the essential distributional properties of each dataset while effectively mitigating technical noise inherent in single-cell measurements. This sampling strategy produces representative profiles that balance the need for noise reduction with preservation of biological signal granularity.

The final stage involves applying SURE again to the aggregated collection of bootstrapped samples, yielding secondary metacells that embody integrated cross-dataset patterns. This hierarchical processing pipeline offers distinct advantages for atlas-scale integration tasks, particularly in terms of computational efficiency and biological fidelity. By enabling parallel processing of individual datasets before final integration, the method scales effectively to handle large cell atlases.

### Public single-cell datasets used in the study

Our evaluation framework utilized multiple publicly available single-cell datasets to assess metacell calling performance across different biological contexts. The primary evaluation employs two carefully selected single-cell datasets: a CD34+ hematopoietic stem and progenitor cell (HSPC)dataset containing approximately 6,800 + sorted cells (available at https://zenodo.org/records/6383269) and a 10x Genomics peripheral blood mononuclear cell (PBMC) dataset comprising 10,000 cells from a single donor (accessible via https://support.10xgenomics.com/single-cell-multiome-atac-gex/datasets/1.0.0/pbmc_granulocyte_sorted_10k). Both datasets were processed following established protocols, with the 10k PBMC dataset annotated according to Muon’s tutorial guidelines.

For cross-batch metacell identification assessment, we incorporated the NeurIPS 2021 single-cell multimodal data integration competition dataset (NCBI GEO accession GSE194122), focusing exclusively on the gene expression modality from this Multiome technology-generated resource. This comprehensive dataset contains over 68,000 cells distributed across 13 batches, providing a robust platform for evaluating batch-consistent metacell identification.

To demonstrate SURE’s capability in atlas augmention through metacell codebooks, we leveraged two lung cell datasets from the CELLxGENE data repository: The Travaglini dataset (accession 5d445965-6f1a-4b68-ba3a-b8f765155d3a) containing over 65,000 lung cells from healthy individuals serves as our reference atlas, while the Chan dataset (accession 62e8f058-9c37-48bc-9200-e767f318a8ec) provides approximately 150,000 cells from both normal tissue and small cell lung carcinoma/lung adenocarnoma patients. These complementary datasets enable rigorous testing of metacell methods across healthy and disease states while maintaining consistent tissue context.

For evaluating large-scale atlas integration capabilities, we utilized three comprehensive CELLxGENE datasets spanning diverse tissues. The DominguezConde dataset (accession 62ef75e4-cbea-454e-a0ce-998ec40223d3) provides extensive coverage with approximately 330,000 cells representing normal human tissues. The Eraslan dataset (accession a3ffde6c-7ad2-498a-903c-d58e732f7470) contributes another 210,000 cells systematically sampled from eight distinct human tissues. For the most comprehensive assessment, we included the TSC dataset (accession e5f58829-1a66-40b5-a624-9046778e74f5), which offers an unprecedented scale with over one million cells encompassing nearly all tissue types from healthy individuals.

For the establishment of HBMCA, a comprehensive resource for studying the human blood mononuclear cell landscape, the atlas datasets utilized, along with their detailed data information and acquisition methods, are provided in the **Table S1**.

To validate the utility and effectiveness of HBMCA, we employed data from 13 different diseases, spanning a wide range of pathological conditions. These datasets were obtained from the latest research articles and are openly accessible in public databases, promoting collaboration and facilitating further investigations by the scientific community. Notably, the data from HBV studies on Asian and European populations can be accessed from the NCBI GEO database under the accession numbers GSE182159 and GSE247322, respectively. In conducting the zero-shot mapping and visualization analysis of single-cell data across multiple diseases, we obtained the datasets from various reputable sources to ensure the reliability and quality of the data. The datasets for dengue virus infection and checkpoint inhibitor-induced colitis were acquired from the NCBI GEO database. These datasets can be accessed using the accession numbers GSE220969 and GSE206298, respectively. To obtain the sepsis dataset, we utilized the Zenodo platform. The dataset can be retrieved using the following DOI: https://doi.org/10.5281/zenodo.7723202. For the acute myocardial infarction data, we accessed the dataset through the eQTLGen Consortium’s website, which can be found at https://eqtlgen.org/sc/datasets/blokland2024.html. The juvenile dermatomyositis dataset was obtained from the CZ CELLxGENE Discover resource, a platform that facilitates the exploration and analysis of single-cell data. The dataset is available at https://cellxgene.cziscience.com/collections/c672834e-c3e3-49cb-81a5-4c844be4a975. To acquire the datasets for multiple sclerosis, myasthenia gravis, systemic lupus erythematosus, and HIV infection, we utilized the Single Cell Portal, a centralized repository for single-cell genomics data. The specific datasets can be accessed using the accession numbers SCP1963 and SCP256. Lastly, the COVID-19 dataset was obtained from the COVID-19 Cell Atlas website (https://www.covid19cellatlas.org/), which serves as a valuable resource for researchers studying the cellular and molecular mechanisms of SARS-CoV-2 infection.

### Comparative analysis of metacell calling methodologies

This study presents a comprehensive comparison between SURE and several state-of-the-art metacell calling approaches, including SEACells, MetaQ, SuperCell, and Metacell2. Each method employs distinct computational strategies for metacell identification, with varying implications for result quality and applicability. SEACells utulizes archetypal analysis to identify metacells that preserve topological structures in latent space, though its current implementation does not directly incorporate transcriptional profiles in the archetypal analysis process, potentially compromising transcriptional homogeneity within metacells. While recent versions have added GPU acceleration support, the core algorithm remains fundamentally designed for smaller datasets.

MetaQ builds upon the VQ-VAE framework, following traditional variational autoencoder architectures that model latent variables using Gaussian distributions. This conventional approach necessitates additional ad hoc operations to regulate metacell sizes, often resulting in suboptimal adjustments that may compromise final metacell quality. SuperCell employs graph-based representation of single-cell data, applying clustering algorithms to detect metacells within the graph structure. Metacell2 represents an evolution of the original Metacell method, implementing a recursive divide-and-conquer algorithm that enables efficient decomposition of scRNA-seq datasets of any size.

### Comparative analysis of atlas integration and query-to-reference mapping

Our study systematically evaluates several advanced atlas-scale integration methods, each implementing distinct computational strategies for single-cell data analysis. SCimilarity employs contrastive learning [45] to optimize cell state integration, effectively clustering similar cellular phenotypes while maintaining separation between distinct populations. It supports zero-shot mapping, allowing direct projection of new datasets onto reference atlases without requiring additional training steps. Scanorama’s specialized panorama stitching algorithm provides unique capabilities for identifying and merging overlapping cellular populations across diverse datasets, with its innovative matching approach specifically designed to handle partial dataset overlaps.

The scVI framework utilizes a variational autoencoder architecture with continuous Gaussian-distributed latent variables, achieving scalability for large-scale atlas data through GPU acceleration. Unlike SURE’s integrated approach, scVI requires supplementary fine-tuning via scArches to enable query-to-reference mapping functionality. Our benchmarking analysis compares the query-to-reference mapping performance of SCimilarity and scArches-enhanced scVI implementations against SURE, providing critical insights into the relative strengths of these different methodological paradigms. This comparative evaluation framework enables rigorous assessment of how various computational strategies address the complex challenges of atlas-scale single-cell integration and query-to-reference mapping.

### Evaluation of metacell calling methods

We utilize mcRigor to assess the performance of SURE and alternative metacell calling methods. The method operates through a rigorous hypothesis testing framework designed to assess metacell homogeneity in single-cell RNA sequencing data. The analysis begins by formalizing the null hypothesis, which states that all cells within a given metacell share an identical multinomial expression profile. This assumption is mathematically expressed as *H*_0_: *x*_*i*_|*θ*∼*Multinomial* (*l*_*i*_, θ_1_, ⋯, θ_*p*_] for every cell *i* within the metacell, where *x*_*i*_ represents the count vector, *l*_*i*_ denotes the library size, and **θ** is the shared probability vector across *p* genes.

The methodology implements a carefully designed permutation strategy to construct appropriate null distributions. The within-feature permutation systematically shufles expression values for each gene independently across all cells in the metacell. For a metacell expression *X* ∈ *R*^*n*×*p*^, this process generates a permuted matrix 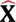 where each column 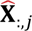 contains randomly reordered values from the original column *X*_:,j_. This approach maintains gene-specific characteristics while effectively nullifying biological interactions.

The method quantifies the deviation of observed correlations from the null expectation through the following ratio:

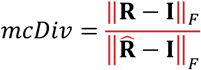

Here, **R** represents the empirical gene-gene correlation derived from actual observations, while 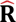 is obtained from the within-feature permuted data. The Frobenius norm serves as the distance metric, measuring the deviation of observed correlations from the identity matrix **I**, which would indicate complete independence between genes. A metacell is identified as potentially dubious when its mcDiv score exceeds a predetermined threshold, suggesting the presence of heterogeneous cell states that violate the homogeneity assumption. Metacells that pass this hypothesis testing are classified as trustworthy.

In our analysis, we use the proportion of trustworthy metacells as the primary metric for evaluating metacell calling methods. mcRigor’s optimization functionality provides a systematical framework for assessing metacell results across varying gamma values.

### Comparison of cell lineage trees

We evaluate the performance of metacell calling methods in preserving biological relationships by analyzing cell lineage trees. For each method under evaluation, we reconstruct a cell lineage tree from its metacell results by using Scanpy’s scanpy.tl.dendrogram function. We also build a cell lineage tree from the original single-cell RNA dataset as the reference. Three distinct but complementary metrics are employed to systematically compare the metacell-derived lineage trees with the reference tree.

The first metric, Kendall’s tau rank correlation coefficient, provides a quantitative measure of leaf node ordering consistency between trees. This nonparametric statistic evaluates the ordinal association between two ranked lists, with its mathematical formulation given by:

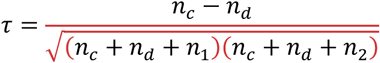

where *n*_*c*_ enumerates concordant pairs that maintain their relative order, *n*_*d*_ counts discordant pairs that exhibit order inversion, and *n*_1_, *n*_2_ account for tied ranks in each respective ordering. The coefficient ranges from −1, indicating perfect inversion of ordering, to 1, representing complete agreement in leaf node arrangement.

For assessing topological similarity at the branching level, we employ Baker’s gamma coefficient. This measure operates by comparing the depth distributions of internal nodes between trees. Given trees *T*_1_ and *T*_2_ with internal node sets *I*_1_ and *I*_2_, we first extract depth vectors:

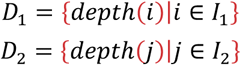

where depth is calculated as the cumulative branch length from root to node. The coefficient is then computed as the Spearman rank correlation between two depth vectors:

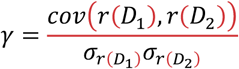

with *r*(⋅) representing the rank transformation, *cov*(⋅,⋅) denoting covariance, and σ indicating standard deviation. This formulation efficiently captures hierarchical similarity in branching patterns while remaining invariant to permulation of leaf nodes.

The third metric, the cophenetic correlation coefficient, evaluates the preservation of pairwise phylogenetic distances between trees. This measure compares two cophenetic distance matrices through Pearson correlation:

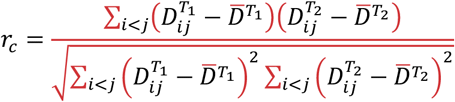

where 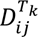 represents the cophenetic distances between leaves *i* and *j* in tree *T*_*k*_, defined as the depth of their lowest common ancestor. The coefficient ranges from 0 to 1, with higher values indicating better preservation of relative cell relationships in the branching structure.

Together, these three metrics form a comprehensive evaluation framework that assesses different but interrelated aspects of tree similarity. Kendall’s tau focuses on the consistency of cell state ordering, Baker’s gamma examines the preservation of branching hierarchy, and the cophenetic correlation evaluates the fidelity of distance relationships between cell states. This orghogonal set of measures provides rigorous quantification of how effectively metacell calling methods maintain the biological hierarchy present in single-cell data.

### Evaluation of atlas-scale integration

We employ scGraph to systematically assess the performance of atlas-scale data integration methods by quantifying how well single-cell embedding preserves biological relationships. This metric operates by comparing cell-type affinity graphs constructed from embeddings against a reference consensus graph constructed from batch-wise PCA loadings, emphasizing the consistency of cell-type neighborhoods across batches while accommodating biologically meaningful variations. Unlike traditional metrics focused solely on batch mixing, scGraph provides complementary insights by evaluating hierarchical and continuous biological structures, providing a more comprehensive assessment of embedding quality.

Additionally, we utilize scIB-metrics, a specialized evaluation framework designed for single-cell genomics, to rigorously benchmark data integration performance. This suite of metrics includes average silhouette width (ASW) and normalized mutual information (NMI) to assess the preservation of distinct cell populations, alongside batch-related metrics such as batch effect removal score (BRAS, an enhanced ASW) and k-nearest-neighbor batch effect test (KBET) to evaluate the effectiveness of batch correction. Together, scGraph and scIB-metrics enable a multi-dimensional evaluation of integration methods.

## Acknowledgements

We thank the members of the Han and Zeng group for their comments and help with preparing the manuscript.

## Authors’ contributions

F.Z. and J.H. conceived the concept and supervised the project. F.Z. designed the model. F.Z. wrote the code. F.Z. performed the experiments. F.Z. and J.H. analyzed the experimental results. F.Z and J.H. drafted the manuscript.

## Funding

This work is supported in part by the Natural Science Foundation of Xiamen, China (No: 3502Z202373020), National Natural Science Foundation of China (82388201, 61503314), Natural Science Foundation of Fujian Province (2019J01041), the National Key R&D Program of China (2020YFA0803500), the CAMS Innovation Fund for Medical Sciences (2019-I2M-5-062).

## Availability of data and materials

Source code of SURE is deposited in Zenodo (DOI: 10.5281/zenodo.16732195) and is freely available for academic use on GitHub at https://github.com/ZengFLab/SURE under the MIT liscense.

## Declarations

### Ethics approval and consent to participate

No ethical approval was required for this study

## Competing interests

The authors declare no competing interests.

## Notes

### Competing Interest Statement

The authors have declared no competing interest.

### Summary of Updates

Rewrite abstract; Perform additional evaluations to systematically compare SURE and alternative methods; Revise Methods to provide better understanding.

